## Supplementary material for "IFNγ and acute prenatal infection with *Toxoplasma gondii* activate fetal hematopoietic stem cells": All Supplement

#### Supplementary Figure Legends

**SFigure 1. Maternal infection with *Toxoplasma gondii* modulates fetal growth and hematopoietic development**

**A)** Visual comparison of crown-rump length from E16.5 fetuses from saline and RH infected mothers, as shown in Fig. 1A.

**B)** Fraction of viable (or "non-resorbed") fetuses per litter observed in E16.5 litters following saline or infection with Pru or RH; n=4 litters/condition.

**C)** Schematic of hematopoietic stem and progenitor (HSPC) cell hierarchy and surface markers.

**D)** Representative gating strategy for fetal liver Tom+ HSCs and GFP+ drHSCs at E16.5.**E-L)** Frequency of **E)** HSPCs **F)** LT-HSCs, **G)** ST-HSCs, **H)** Tom+ HSCs, **I)** GFP+ drHSCs, **J)** MPP2, **K)** MPP3, **L)** MPP4 in E16.5 fetuses following saline or maternal infection with Pru or RH as shown in Fig. 1A. n = 9-15 fetuses from 3 litters per each condition.

**M)** Total cellularity of E16.5 fetal liver cells following saline or infection with Pru or RH.

**N)** Total cellularity of CD45+ cells in fetal liver at E16.5 following saline or infection with Pru or RH. For E and N, each dot represents results from an individual fetus, 4 litters were analyzed per each condition. For all analysis above bars represent mean. Data analyzed by one-way ANOVA with Tukey's test. $^*p \leq 0.05$; $^{**}p \leq 0.01$; $^{***}p \leq 0.001$; $^{****}p \leq 0.0001$.

**SFigure 2. Maternal infection with *Toxoplasma gondii* leads to lasting changes in BM reconstitution of fetal HSCs.**

**A-M)** Bone Marrow (BM) chimerism of **A)** HSPCs, **B)** LT-HSCs, **C)** ST-HSCs, **D)** MPP2, **E)** MPP3, **F)** MPP4, **G)** granulocyte macrophage progenitors (GMP), **H)** megakaryocyte progenitors (MkP), **I)** erythroid progenitors (EP), **J)** granulocyte/macrophages (GM), **K)** common lymphoid progenitors (CLP), **L)** B-cells, and **M)** T-cells in primary recipients of Tom+ HSCs or GFP+ drHSCs at 18-weeks post-transplantation. n of mice is shown as the numerator in Fig. 2B.

**N-Z)** Bone Marrow (BM) chimerism of **N)** HSPCs, **O)** LT-HSCs, **P)** ST-HSCs, **Q)** MPP2, **R)** MPP3, **S)** MPP4, **T)** granulocyte macrophage progenitors (GMP), **U)** megakaryocyte progenitors (MkP), **V)** erythroid progenitors (EP), **W)** granulocyte/macrophages (GM), **X)** common lymphoid progenitors (CLP), **Y)** B-cells, and **Z)** T-cells in secondary Tom+ HSC or GFP+ drHSC transplant recipients at 18-weeks post-transplant. n of mice is shown as the numerator in Fig 2K. For all analyses above, bars represent mean $\pm$ SD. Statistical significance was determined by one-way

ANOVA with Tukey's test. *p ≤ 0.05; **p ≤ 0.01; ***p ≤ 0.001. No reconstitution was present for the secondary recipients of the GFP+ drHSC under RH condition.

**SFigure 3: Bone marrow chimerism from IFNγ exposed fetal HSCs following transplantation.**

**A-M)** Bone Marrow (BM) chimerism of **A)** HSPCs, **B)** LT-HSCs, **C)** ST-HSCs, **D)** MPP2, **E)** MPP3, **F)** MPP4, **G)** granulocyte macrophage progenitors (GMP), **H)** megakaryocyte progenitors (MkP), **I)** erythroid progenitors (EP), **J)** granulocyte/macrophages (GM), **K)** common lymphoid progenitors (CLP), **L)** B-cells, and **M)** T-cells in primary recipients of Tom+ HSCs or GFP+ drHSCs at 18-weeks post-transplantation. N of mice is shown as the numerator in Fig. 5A.

**N-Z)** Bone Marrow (BM) chimerism of **N)** HSPCs, **O)** LT-HSCs, **P)** ST-HSCs, **Q)** MPP2, **R)** MPP3, **S)** MPP4, **T)** granulocyte macrophage progenitors (GMP), **U)** megakaryocyte progenitors (MkP), **V)** erythroid progenitors (EP), **W)** granulocyte/macrophages (GM), **X)** common lymphoid progenitors (CLP), **Y)** B-cells, and **Z)** T-cells in secondary recipients of Tom+ HSC or GFP+ drHSCs at 18-weeks post-transplantation. N of mice is shown as the numerator in Fig. 5J. For all experiments above, bars represent mean $\pm$ SD. Statistical significance was determined by one-way ANOVA with Tukey's test. *p ≤ 0.05; **p ≤ 0.01; ***p $\leq$ 0.001.

**SFigure 4: The role of IFNγ and IFNγR in fetal response**

**A)** Comparison of IFNγ cytokine in E15.5 fetal amniotic fluid between IFNγR +/- and -/- pups from either IFNγR +/- or -/- dams following saline injection.

**B)** Comparison of IFNγ cytokine in E15.5 fetal amniotic fluid between IFNγR +/- and -/- pups from either IFNγR +/- or -/- dams following IFNγ injection.

**C)** Comparison of IFNγ cytokine in E15.5 fetal liver supernatant between IFNγR +/- and -/- pups from either IFNγR +/- or -/- dams following saline injection.

**D)** Comparison of IFNγ cytokine in E15.5 fetal liver supernatant between IFNγR +/- and -/- pups from either IFNγR +/- or -/- dams following IFNγ injection.

**E)** Comparison of IFNγ cytokine in E15.5 maternal serum between IFNγR +/- and -/- dams following saline or IFNγ injection.

**F)** Population changes in HSCs and MPPs in IFNγR +/- and -/- pups from either IFNγR +/- or -/- dams following IFNγ injection.

For amniotic fluid and fetal liver supernatant, n = 7-10 fetuses from 2-4 litters per condition. For maternal serum, n = 2-4 dams per condition. For all experiments above, bars represent mean $\pm$ SD. Statistical significance was determined by 2-way ANOVA. *$p \leq 0.05$; **$p \leq 0.01$; ***$p \leq 0.001$; ****$p \leq 0.0001$. N=10-14 fetuses from at least 3 litters/condition. ó

**SFigure 1: Inflammation from maternal infection with *Toxoplasma gondii* modulates fetal growth and hematopoietic development**

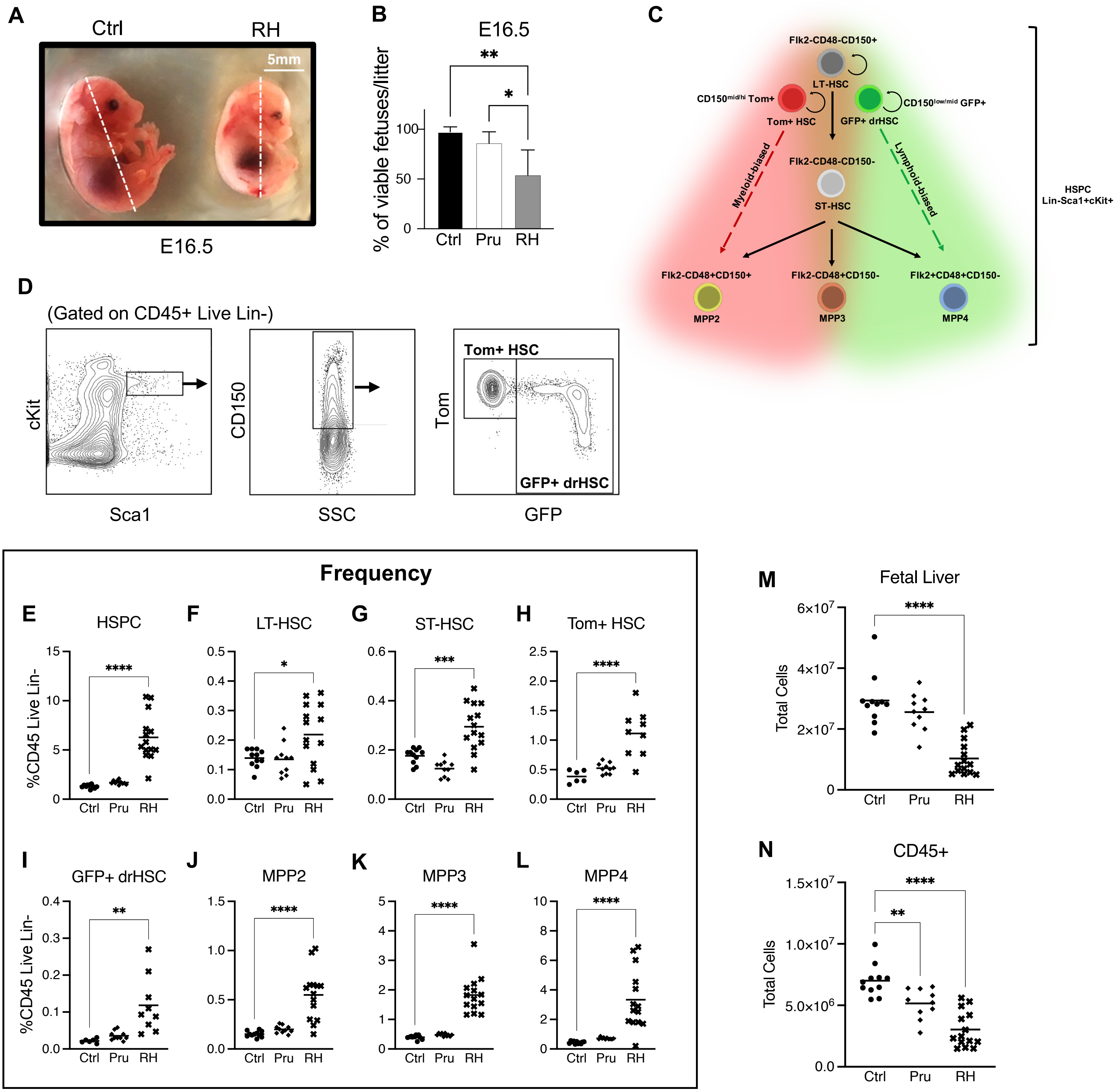

### SFigure 2: Maternal infection with Toxoplasma gondii leads to lasting changes in BM reconstitution of fetal HSCs

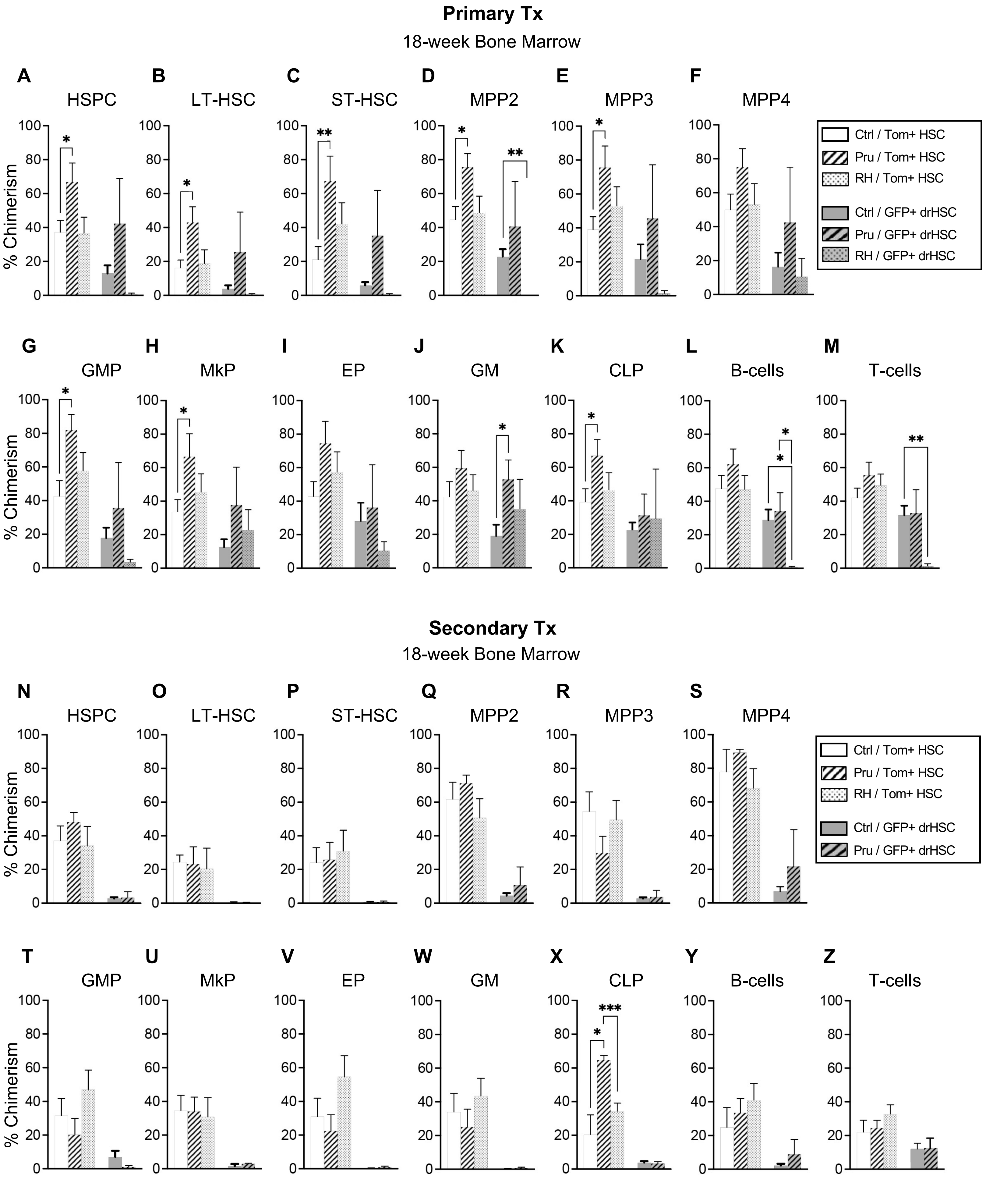

### SFigure 3: Bone marrow chimerism from IFNγ exposed fetal HSCs following transplantation

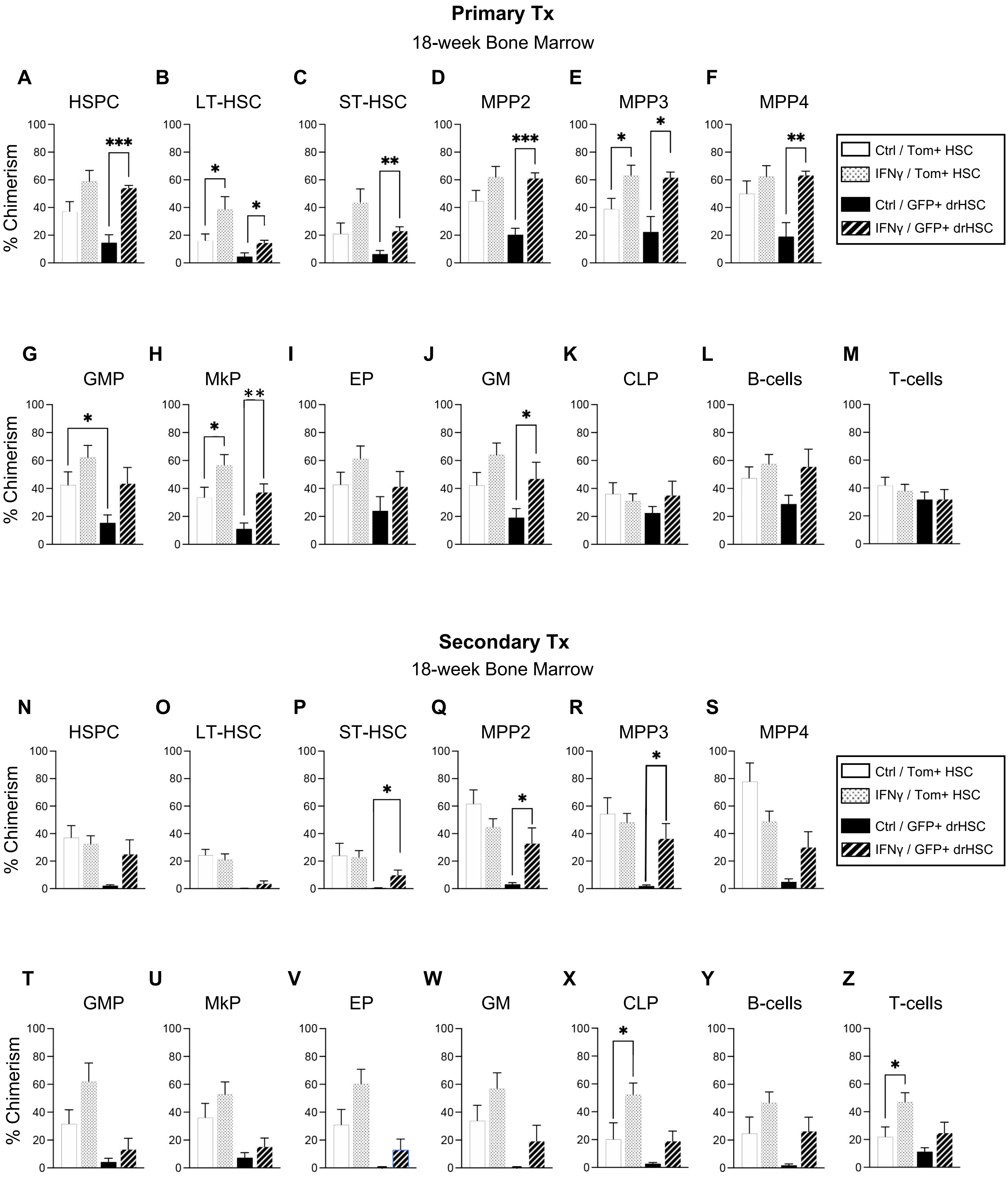

#### SFigure 4: The role of IFNγ and IFNγR in fetal response

##### Amniotic Fluid

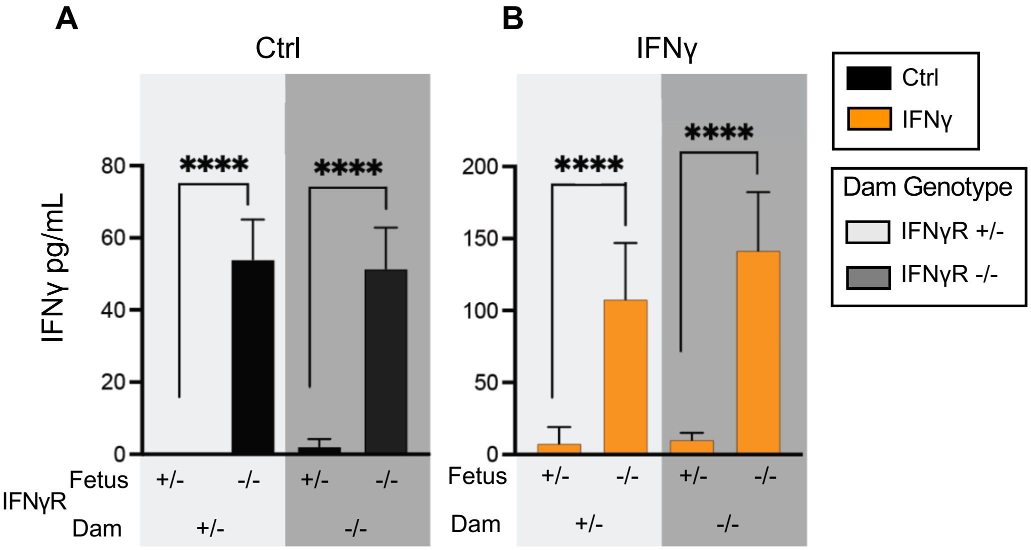


##### Fetal Liver

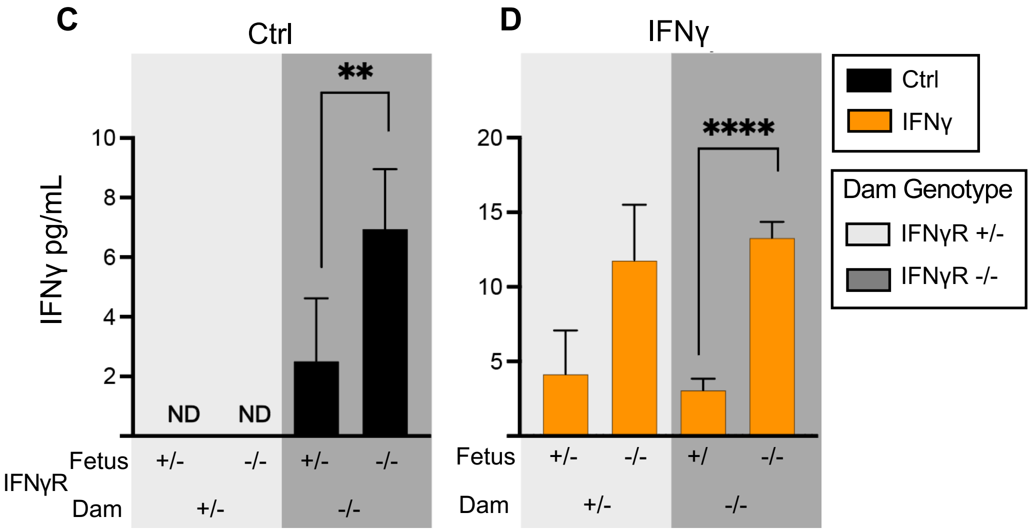


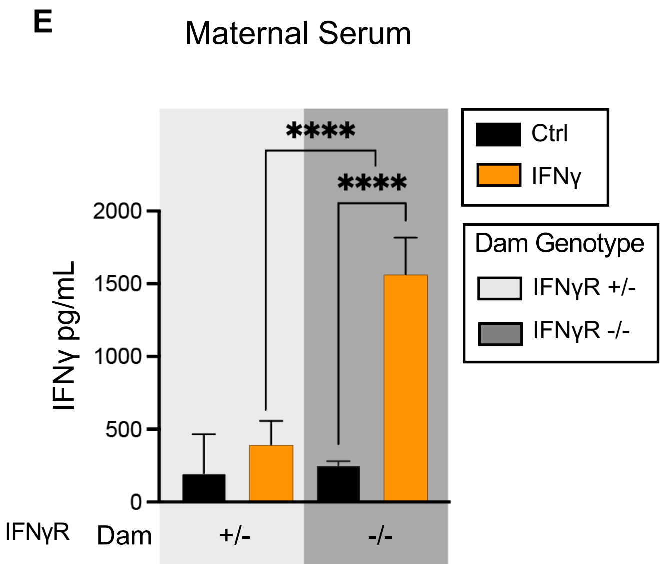


**F**

| **IFNγR** | | **HSCs** | **MPPs** |
|---|---|---|---|
| **Dam** | **Fetus** | | |
| +/- | +/- | Up | Up |
| +/- | -/- | No change | Up |
| -/- | +/- | No change | Up |
| -/- | -/- | No change | No change |
